# Homologous recombination changes the context of *Cytochrome b* transcription in the mitochondrial genome of *Silene vulgaris* KRA

**DOI:** 10.1101/422808

**Authors:** Helena Štorchová, James D. Stone, Daniel B. Sloan, Oushadee Abeyawardana, Karel Müller, Jana Walterová, Marie Pažoutová

**Affiliations:** Institute of Experimental Botany v.v.i, Academy of Sciences of the Czech Republic, Plant Reproduction Laboratory, Rozvojová 263, Prague, 16502 Czech Republic; Colorado State University, Department of Biology, Fort Collins, CO 80523, USA

## Abstract

**Background:** *Silene vulgaris* (bladder campion) is a gynodioecious species existing as two genders – male-sterile females and hermaphrodites. Cytoplasmic male sterility (CMS) is generally encoded by mitochondrial genes, which interact with nuclear fertility restorer genes. Mitochondrial genomes of this species vary in DNA sequence, gene order and gene content. Multiple CMS genes are expected to exist in *S. vulgaris*, but little is known about their molecular identity.

**Results:** We assembled the complete mitochondrial genome from the haplotype KRA of *S. vulgaris*. It consists of five chromosomes, two of which recombine with each other. Two small non-recombining chromosomes exist in linear, supercoiled and relaxed circle forms. We compared the mitochondrial transcriptomes from females and hermaphrodites and confirmed the differentially expressed chimeric gene *bobt* as the strongest CMS candidate gene in *S. vulgaris* KRA. The chimeric gene *bobt* is co-transcribed with the *Cytochrome b* (*cob*) gene in some genomic configurations. The co-transcription of a CMS factor with an essential gene may constrain transcription inhibition as a mechanism for fertility restoration because of the need to maintain appropriate production of the necessary protein. Homologous recombination places the gene *cob* outside the control of *bobt*, which allows for the suppression the CMS gene by the fertility restorer genes. In addition, by analyzing RNA editing, we found the loss of three editing sites in the KRA mitochondrial genome and identified four sites with highly distinct editing rates between KRA and another *S. vulgaris* haplotypes (KOV). Three of these highly differentially edited sites were located in the *transport membrane protein B* (*mttB*) gene. They resulted in differences in MttB protein sequences between haplotypes despite completely identical gene sequences.

**Conclusions:** Frequent homologous recombination events that are widespread in plant mitochondrial genomes may change chromosomal configurations and also the control of gene transcription including CMS gene expression. Posttranscriptional processes, e.g RNA editing shall be evaluated in evolutionary and co-evolutionary studies of mitochondrial genes, because they may change protein composition despite the sequence identity of the respective genes. The investigation of natural populations of wild species such as *S. vulgaris* are necessary to reveal important aspects of CMS missed in domesticated crops, the traditional focus of the CMS studies.

## Introduction

Flowering plants possess two organelles with genomic DNA - plastids and mitochondria. Mitochondrial (mt) genomes are more variable than plastid genomes in both size and structure. MtDNA ranges in size from 66 kb (hemiparasitic mistletoe *Viscum scurruloideum*, [1] up to 11 Mb (*Silene conica*, [2]. Intergenic regions of unknown origin and DNA transferred from the nucleus or plastids are mainly responsible for this enormous size variation [3], whereas the number of genes varies by only about two-fold across angiosperms. Most angiosperm mt genomes contain 24 to 41 protein coding genes, three genes for rRNA and a variable number of tRNA genes ([4, 5]. Gene order is not conserved, and a general lack of synteny has been documented even at the intraspecific level ([6, 7, 8, 9].

Frequent rearrangements of plant mt genomes are primarily caused by intramolecular recombination. Homologous recombination across large repeats (> 500 bp) is common and leads to an equilibrium among alternative genomic configurations ([10] Marechal and Brisson 2010). Recombination between shorter repeats is less frequent and may result in changes in stoichiometry of individual molecular variants [11, 12, 13]. Microhomology-mediated recombination across very short repeats < 50 bp [14] is rare and may generate chimeric genes composed of fragments of mt genes and intergenic regions. The frequency of microhomology-mediated recombination is increased by mutations in the nuclear genes responsible for recombination surveillance such as *MUTS HOMOLOG 1* (*MSH1*) [15] or *HOMOLOG OF BACTERIAL RECA 3* (*RECA3*) [16].

Homologous recombination in plant mtDNA participates in the repair of double-strand breaks [17, 18], which arise due to DNA damage [7]. Respiration in plant mitochondria requires electron transfer through the electron transport chain located in the inner membrane. The electron transport chain can also lead to the production of reactive oxygen species (ROS) such as peroxides, superoxides, and hydroxyl radicals, which damage mtDNA [19]. Whereas homologous recombination is associated with accurate DNA repair, alternative RECA-independent pathways e.g. break-induced replication are involved in error-prone repair, which generates major structural rearrangements and new repeats, which in turn serve as a substrate for additional recombination [18, 20, 21].

Unlike the fast and dynamic structural evolution of plant mt genomes, substitution rates in mt genes are generally slow [22, 23, 24]. However, some phylogenetic lineages exhibit accelerated mt substitution rate, including the genus *Silene* [25]. Substitution rates in this genus vary >100-fold across species [2, 26].

*Silene vulgaris* (bladder campion) shows only a slightly elevated mt substitution rate, but it exhibits high levels of within-species polymorphism in mtDNA, with variation not only in gene order and intergenic sequences, but also gene content in these already highly reduced genomes [9]. The completely sequenced mt genomes of four *S. vulgaris* haplotypes are multichromosomal and highly rearranged. They contain numerous repeats of various sizes undergoing frequent recombination and generating chimeric open reading frames (ORFs), which may cause cytoplasmic male sterility (CMS) [9].

CMS is an example of a cyto-nuclear interaction, which results in the production of male sterile (female) and hermaphroditic plants [27]. Mitochondrial-encoded CMS genes (often but not always chimeric ORFs) interfere with mt metabolism and prevent the development of viable pollen [28, 29]. Their expression is inhibited by nuclear fertility-restorer (*Rf*) genes, which re-establish male fertility in hermaphroditic plants [30]. The reproduction system termed gynodioecy is often caused by CMS. It is characterized by the co-occurrence of females and hermaphrodites in the same population and is widespread among flowering plants [31]. Mitochondrial transmission in females of gynodioecious species may be augmented relative to hermaphrodites owing to exclusive resource allocation to ovules and avoidance of inbreeding depression [32, 33]. Models suggest that CMS genes can spread in the population if they are rare. When their frequency increases, pollen limitation may select for an increase of matching *Rf* genes, eliminating the selective advantage of the CMS gene. The whole scenario may repeat when a different CMS gene requiring distinct *Rf* invades the population [32]. Such processes in individual populations have been hypothesized to establish balancing selection through negative frequency dependence, which maintains polymorphism of organellar genomes at metapopulation level [32, 34].

Sequence variation in plastid and mt loci is often higher in gynodioecious species than in their dioecious or hermaphroditic congeners, which could indicate the action of balancing selection [35, 36, 37]. The comparative study of *Silene nutans* and *Silene otites* by [38] found a higher cytoplasmic diversity in gynodioecious *S. nutans* than in dioecious *S. otites* despite a faster mt substitution rate in the latter. High polymorphism of mtDNA in *S. vulgaris* [39, 40, 41] may, therefore, be related to its gynodioecious mating system.

*S. vulgaris* has been investigated in multiple population genetic studies [41, 42, 43, 44], but a detailed understanding of the evolutionary processes responsible for its extremely rearranged and diverse mt genomes [9] is still lacking. The comprehensive transcriptomic study of the mt haplotype KOV of *S. vulgaris* lacking chimeric ORFs revealed a mt long non-coding RNA associated with CMS [45]. This finding suggests that CMS types in this species are very diverse. Detailed investigation of various CMS types associated with particular mt haplotypes of *S. vulgaris* may shed light on the complex processes shaping mt genomes in gynodioecious plants.

Here, we report the mt genome and transcriptome of an *S. vulgaris* accession collected near Krasnoyarsk (Siberia, Russia) to expand the geographical sampling to Asia and to get detailed information about the mt haplotype, where the CMS candidate gene *bobt* was identified previously [46]. We confirmed the chimeric *bobt* gene to be the most likely CMS factor. We also discovered that homologous recombination can move *bobt* immediately upstream of the essential gene for cytochrome b (*cob*), leading to their co-transcription. The ratio of the two configurations - *cob* under the control of *bobt* or under the control of its typical promoter – varied among plants. In addition, we found that C-to-U RNA editing differed between the mt haplotypes KOV and KRA. We identified three independent losses of editing sites, which implies that the loss of editing sites can occur rapidly. These findings provide insights into the dynamics of mt genomes in natural populations with CMS.

## Results

### Mitochondrial genome of S. vulgaris KRA

We obtained 76 Mb of DNA sequence (343,000 paired-end reads; average fragment length 3 kb) from 454 sequencing run of enriched mt DNA from *S. vulgaris* KRA, which provided > 80× coverage of the mt genome. The assembled mt genome consists of five chromosomes, three of which lack any identifiable repeats that would allow recombination to merge them into a larger “master circle” conformation. Therefore, these chromosomes appear to be “autonomous” from the rest of the genome. The other two may be joined together by homologous recombination (Table 1, Fig. 1). The size of the KRA genome (404,739 bp) falls within the range of four other mt genomes of *S. vulgaris* that were published previously [9]. The same is true for KRA genome complexity (388 kb), which is the amount of unique sequence after the exclusion of duplicated regions. Eleven repeats > 500 bp and 120 repeats > 100 bp are responsible for the very dynamic character of the KRA genome. The provided circular maps (Additional file 1: Figure S1), therefore, represent only one of many alternative genomic configurations coexisting within the mitochondria of *S. vulgaris* KRA. Genic content (26 protein coding genes, 4 tRNA and 3 rRNA genes) account for 12.5% of the genome. A small portion (9,875 bp; 2.4%) of the KRA genome was derived from plastid DNA. Only about one third of this (3,675 bp) is unique to KRA, the remaining plastid inserts are shared with other *S. vulgaris* mt genomes [9], which suggests their transfer from the plastids before the mt haplotypes diverged.

**Table 1.** Characteristics of the mt genome of *S. vulgaris* KRA.

|  | **Chr 1** | **Chr 2** | **Chr 3** | **Chr 4** | **Chr 5** |
| --- | --- | --- | --- | --- | --- |
| Type | Main | Non-autonomous | Autonomous | Autonomous | Autonomous |
| Length (bp) | 37 305 | 18 063 | 9 488 | 2 578 | 1 558 |
| G + C content (%) | 41.7 | 41.7 | 43.8 | 47.8 | 45.9 |
| Intact protein genes<sup>a</sup> | 23 | 2 | 1 | 0 | 0 |
| tRNA genes | 4 | 0 | 0 | 0 | 0 |
| rRNA genes | 3 | 0 | 0 | 0 | 0 |
| GenBank accession | MH455602 | MH455603 | MH455604 | MH455605 | MH455606 |

<sup>a</sup> Gene counts include fragments that are *trans*-spliced with exons on other chromosomes to form an intact transcript, each gene was counted only once.

**Figure 1.**
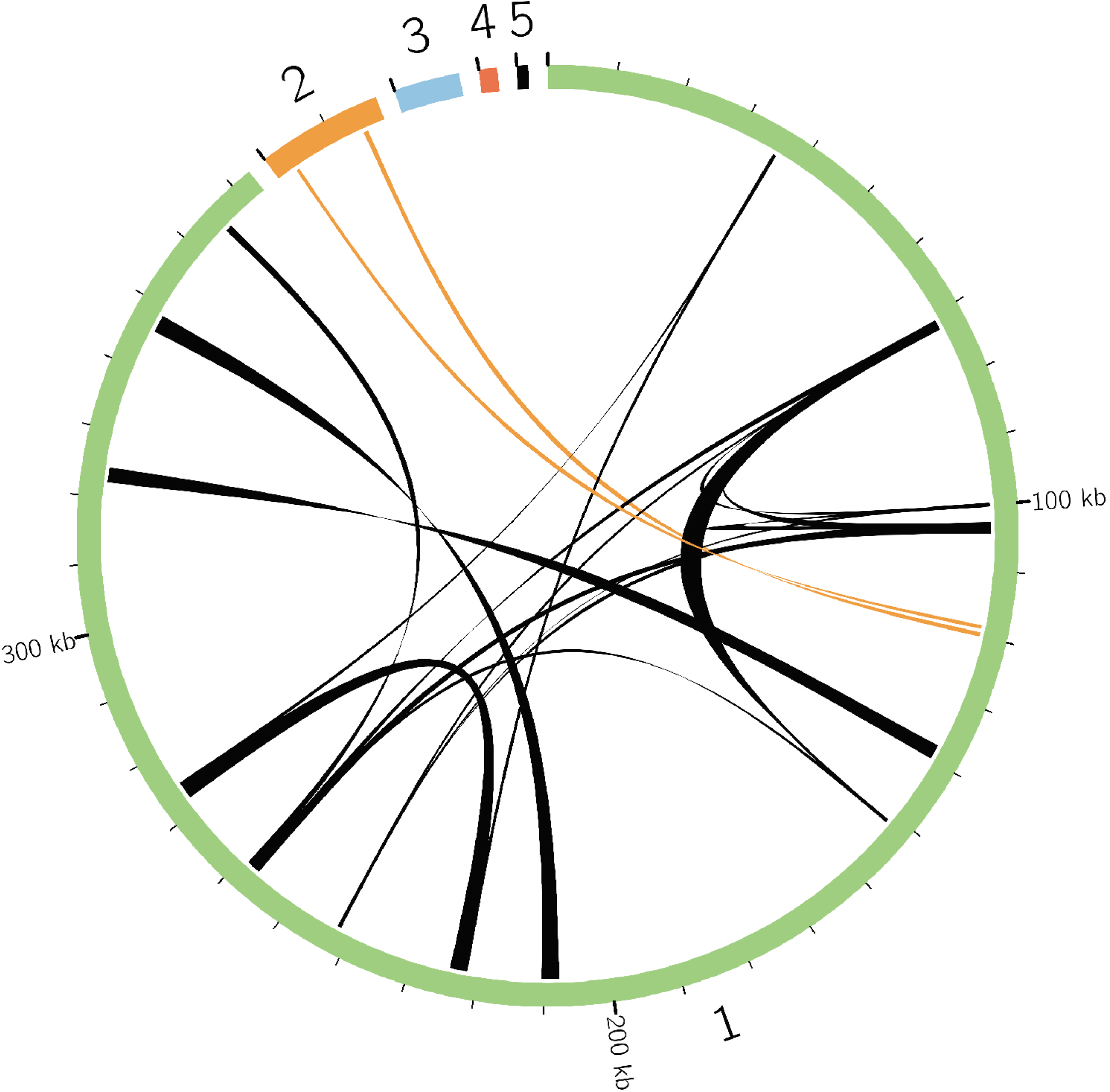
Schematic representation of five chromosome of the mt genome *S. vulgaris* KRA. Homologous recombination across the repeats > 400 bp is depicted by the ribbons. Orange ribbons show the recombination between chromosome 1 and 2.

The KRA mt genome most closely resembles the S9L genome, with which it shares more than 70% of sequence (Table 2). However, about 14% of DNA is not similar to any known sequence (even those from other *S. vulgaris* haplotypes), which illustrates the extraordinary genetic variation in mt genomes within this species. The phylogenetic tree constructed from the concatenated sequences of mt protein genes confirms the relatedness of the KRA and S9L mt haplotypes (Additional file 2; Figure S2). The *atp6* gene in S9L is highly divergent from other haplotypes of *S. vulgaris*, which have the same ancestral *atp6* sequence as *S. latifolia* [9]. The KRA *atp6* haplotype does not share the single nucleotide polymorphisms (SNPs) with S9L, but it contains five unique polymorphic sites, two of which are non-synonymous, and lacks several polymorphic sites unique to KOV (Additional file 3: Table S1).

**Table 2.** Shared sequence content between pairs of *S. vulgaris* mt genomes.

|  | KOV (%) | MTV (%) | S9L (%) | SD2 (%) | KRA (%) | Any (%) | All (%) |
| --- | --- | --- | --- | --- | --- | --- | --- |
| KOV | - | 54.2 | 56.4 | 53.1 | 50.9 | 74.9 | 31.1 |
| MTV | 44.2 | - | 59.8 | 58.2 | 55.3 | 82.9 | 25.7 |
| S9L | 47.1 | 61.1 | - | 54.9 | 73.2 | 91.2 | 26.6 |
| SD2 | 44.3 | 58.2 | 54.1 | - | 54.6 | 81.7 | 26.0 |
| KRA | 42.9 | 55.5 | 71.3 | 54 | - | 85.8 | 26.6 |

The KRA and S9L mt genomes share homologous autonomous chromosomes. Chromosome 3 in the KRA genome and chromosome 6 in the S9L genome are nearly identical (similarity 99.94%) and contain the same genes. Autonomous chromosome 4 of the KRA genome (size 2,578 bp) is similar to the smallest chromosomes in the other *S. vulgaris* genomes [9] (Additional file 4: Figure S3). It contains no genes, but it shares a short region of occasional transcription with KOV chromosome 6 [45]. Sequence similarity of the shared region across five completely sequenced mt genomes of *S. vulgaris* is 96% – 98%, when only nucleotide substitutions, not indels, were taken into account. A putative RNA hairpin structure is located within this area.

### Structure of two small autonomous chromosomes in the KRA mt genome

All four previously sequenced *S. vulgaris* mt genomes [9] share a small autonomous chromosome with similar nucleotide sequence to chromosome 4 in the KRA mt genome. However, unlike in these other haplotypes, this chromosome is not the smallest one in the mt KRA genome. Chromosome 5 is sized only 1,558 bp. It harbors no genes and the majority of its sequence shows no similarity to any GenBank record. A repeat of 287 nt is shared with chromosome 1 (94% similarity), but no recombinant sequence reads were identified.

We employed Southern hybridization to gain insight into the structure of the small chromosomes. When total DNA extracted from leaves or flower buds was digested with *Bgl*II, cutting both chromosomes only once, the fragments corresponding to the expected sizes of linearized chromosomes were obtained (Fig. 2, 3). No additional fragments recombinant confirmations were observed, except for a very faint band matching the size of a linear dimer of the chromosome 4, which could have arisen from an incomplete digestion (Fig. 2).

**Figure 2.**
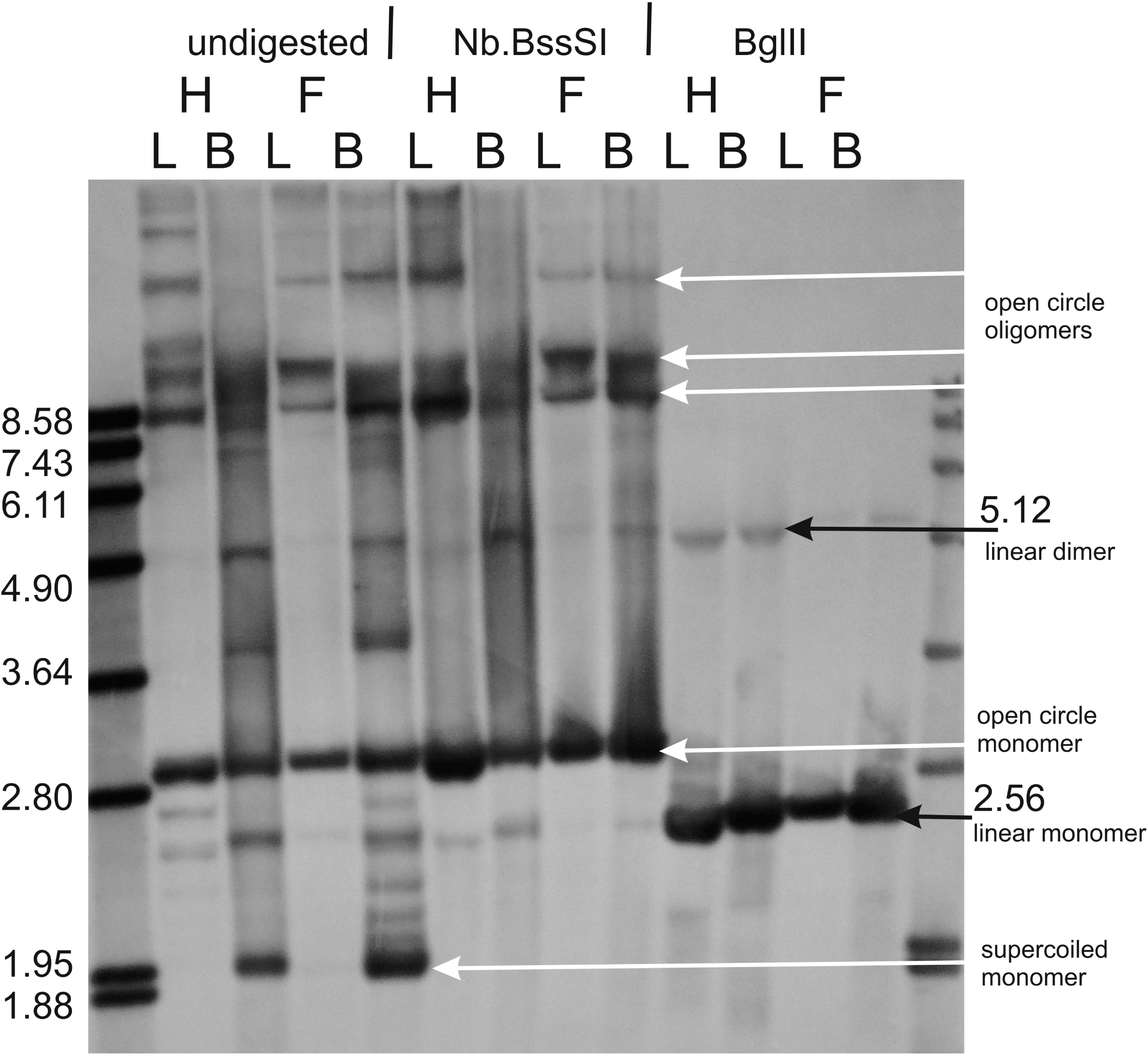
Structure of the autonomous chromosome 4 of *S. vulgaris* KRA. Southern blot hybridization probed with 626 bp long sequence specific to chromosome 4. Total DNAs were extracted from leaf (L) or flower bud (B) tissues of two female (F) and two hermaphroditic (H) plants. DNAs were either digested with the restriction enzyme *Bgl*II recognizing a single site in chromosome 4 (right), treated with nickase *Nb.Bss*SI capable of introducing one single-strand cut in chromosome 4 (middle), or not digested at all (left). Molecular weight standards were loaded on both sides of the 1% agarose gel. Black arrows point to the bands corresponding to linear monomer (2.56 kb) or dimer (5.12 kb) of chromosome 4. White arrows point to the supercoiled monomer or open (relaxed) circle monomer or oligomers.

**Figure 3.**
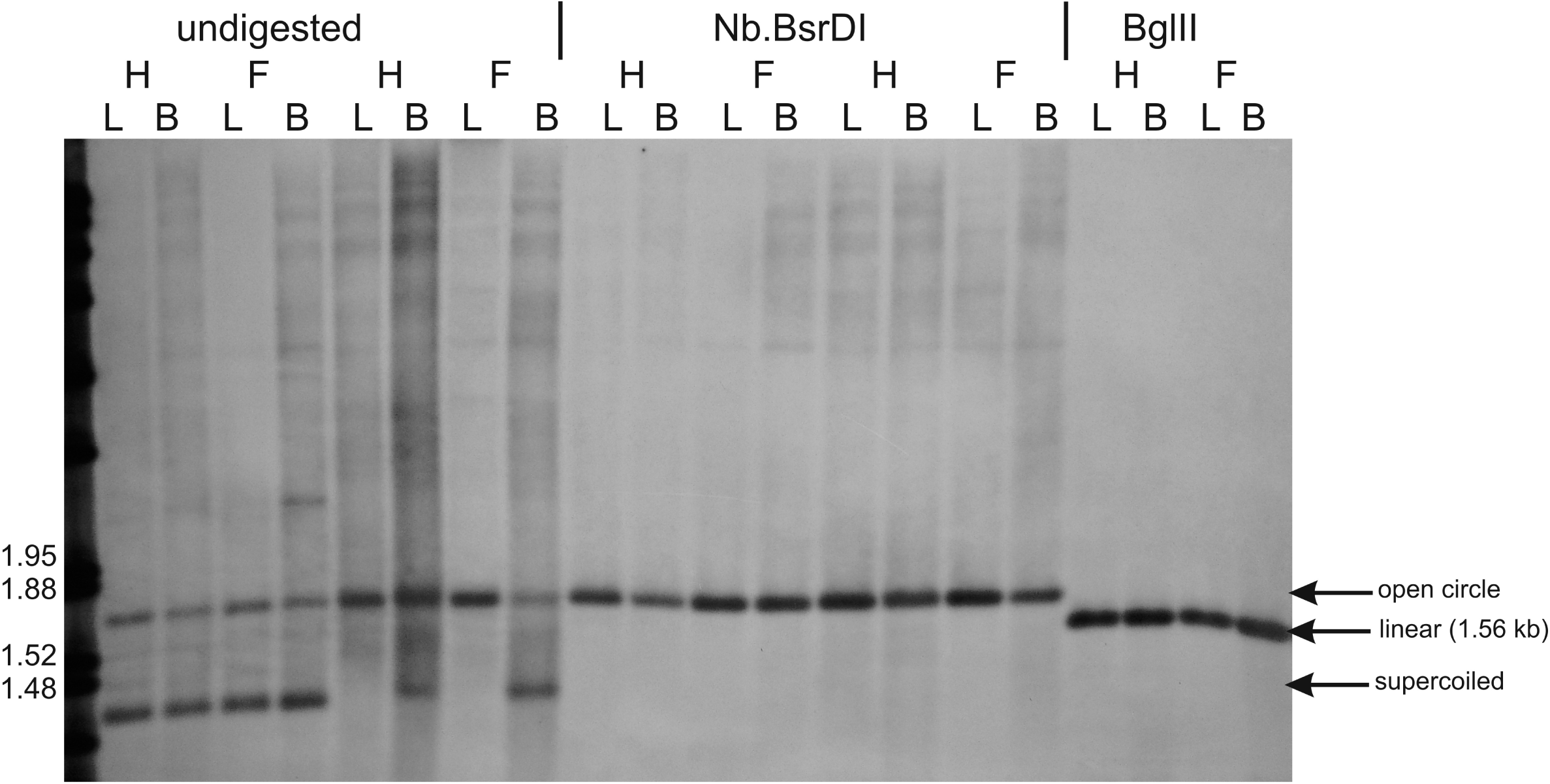
Structure of the autonomous chromosome 5 of *S. vulgaris* KRA. Southern blot hybridization probed with 626 bp long sequence specific to the chromosome 5. Total DNAs were extracted from leaf (L) or flower bud (B) tissues of female (F) and hermaphroditic (H) plants. DNAs were digested with the restriction enzyme *Bgl*II recognizing a single site in chromosome 5 (right), treated with nickase *Nb*.*Bsr*D1 capable of introducing one single-strand cut in chromosome 5 (middle), or not digested at all (left). The arrows point at the bands corresponding to open (relaxed) circle, linear (1.56), or supercoiled monomer of chromosome 5.

Undigested DNA hybridized with a chromosome 4-specific probe provided the most complex banding pattern. To distinguish supercoiled and relaxed (open) circle structures, we digested DNA with nickase Nb.*Bss*SI, which is predicted to generate a single-strand break in one position of chromosome 4. The fastest migrating band and some additional bands were not present after the Nb.*Bss*SI treatment, which implied their supercoiled structure. The remaining bands may correspond to relaxed forms of various oligomers. Interestingly, the banding pattern of undigested DNA extracted from leaves resembled the pattern obtained after digestion with nickase (Fig. 2). The differences between banding patterns of undigested DNA from leaves and flower buds (from which the previous samples were derived) are therefore caused by a different proportion of supercoiled molecules between the two organs.

DNA hybridized with the probe specific to a unique region of chromosome 5 generated a less complicated pattern (Fig. 3). Only one band, corresponding to the relaxed monomer, was detected after the treatment with nickase Nb.*Bsr*DI. The three bands observed in the undigested DNA may be therefore interpreted as relaxed, linear, and supercoiled monomers, similar to the general structure of bacterial plasmids. We observed the differences in the content of the supercoiled form between leaves and flower buds, but not in all plants. Unlike chromosome 4, no band corresponding to oligomers of chromosome 5 was found. No differences in banding patterns between female (F) and hermaphroditic (H) plants were observed by any treatment.

We applied qPCR to estimate the copy number of chromosome 4 and chromosome 5 relative to *rrn18*, which is present in a single copy on the main chromosome 1 (Fig. 4a). The measurements indicated that the small chromosomes existed in several copies for each copy of chromosome 1. The relative copy numbers of chromosome 4 and 5 were significantly lower in floral buds than in leaves (ANOVA, p < 0.023 for chromosome 4, p < 0.000001 for chromosome 5). No significant differences were found between the genders. In contrast, the copy numbers of chromosomes 4 and 5 relative to nuclear rDNA were similar in leaves and flower buds (Fig. 4b).

**Figure 4.**
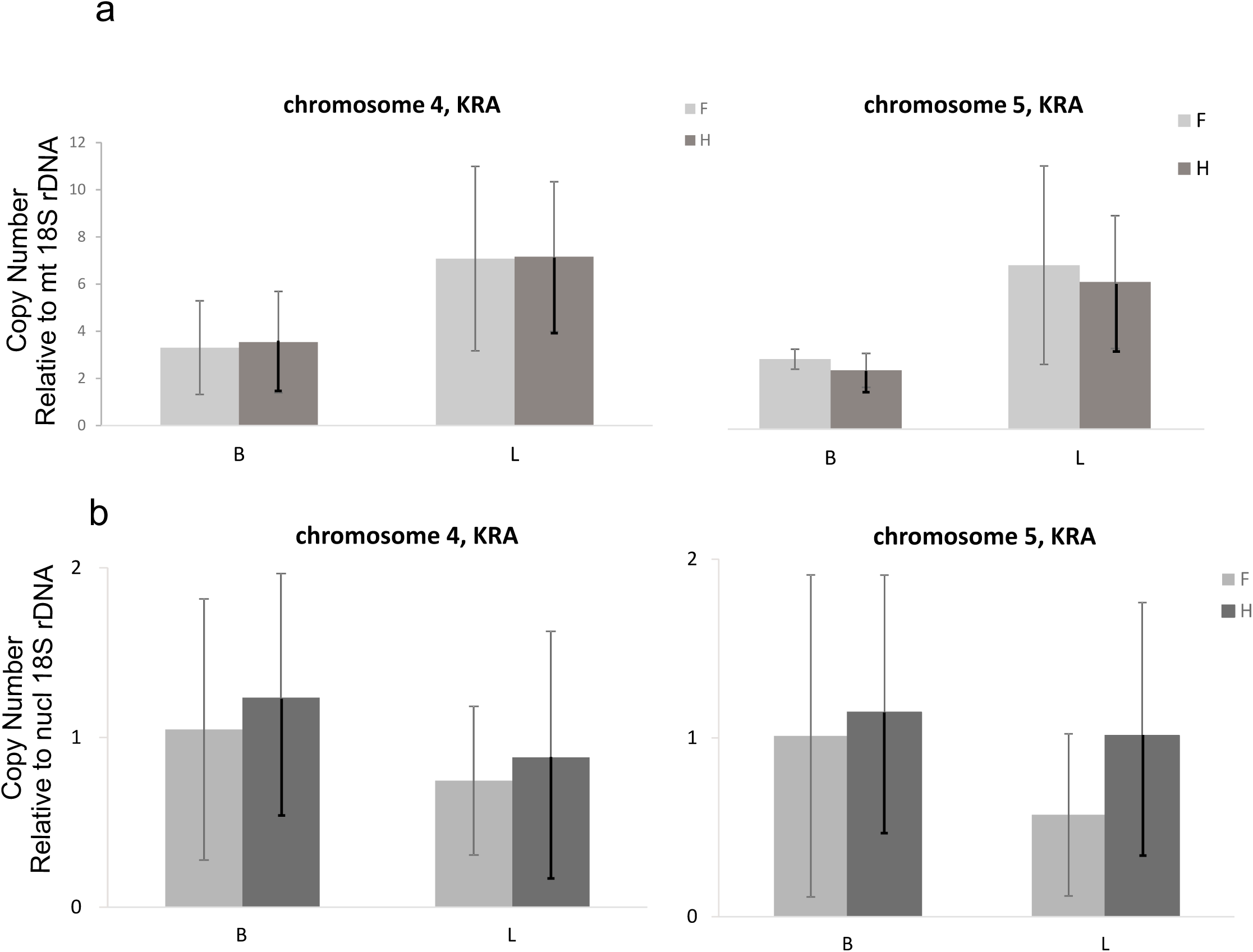
Copy numbers of chromosome 4 and chromosome 5 of the KRA mt genome. Copy numbers were estimated with qPCR relative to mt 18S rDNA (a), or relative to nuclear 18S rDNA (b).

### Homologous recombination across cob and atp6 repeats

The outcome of intramolecular homologous recombination depends on the orientation of repeats. Recombination across direct repeats results in excision of the inter-repeat region and generation of a separate DNA molecule, whereas recombination across inverted repeats leads to the inversion of the inter-repeat region (Fig. 5).

**Figure 5.**
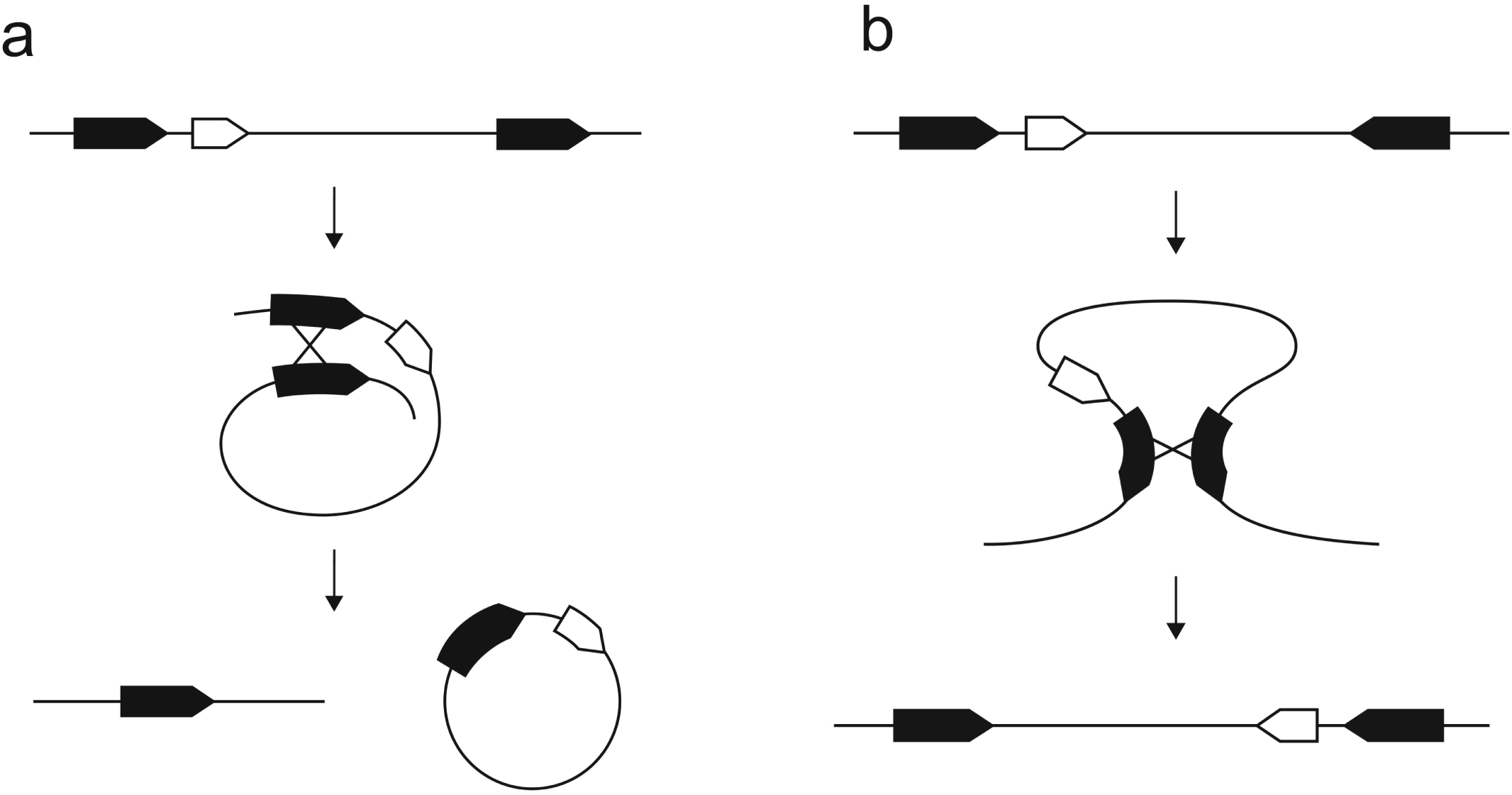
Homologous recombination across direct and inverted repeats. Homologous recombination across direct repeats results in the formation of a separate circular DNA molecule (a), homologous recombination across inverted repeats results in the inversion of the inter-repeat region (b).

The complete *cob* and *atp6* genes are found on chromosome 2, whereas a chimeric ORF (343 amino acids) containing partial sequences of *cob* located at position 118,360 on chromosome 1 (1.118360(orf)(343)) and another chimeric ORF (230 amino acids) containing a portion of *atp6* also residing on chromosome 1 (1.119388(orf)(230)). Recombination across the 636 bp long *atp6* repeat, or across the repeat comprised of 454 bp of *cob* coding sequence and 37 bp of sequence upstream of *cob*, can join these sequences from chromosomes 1 and 2 together. The joint chromosome contains the two repeat pairs in alternating configuration – the *atp6* repeat is placed next to the *cob* repeat (Fig. 6a). Homologous recombination across *atp6* (Fig. 6b) changes the orientation of the *cob* repeats and *vice versa* (Fig. 6c). When inverted repeats gain the direct orientation, additional recombination across them generates a separate DNA molecule, and chromosome 2 is re-established (Fig. 6d, e). All of these predictions about the effects of recombination on alternative genome rearrangements have been confirmed by Southern blots (see below).

**Figure 6.**
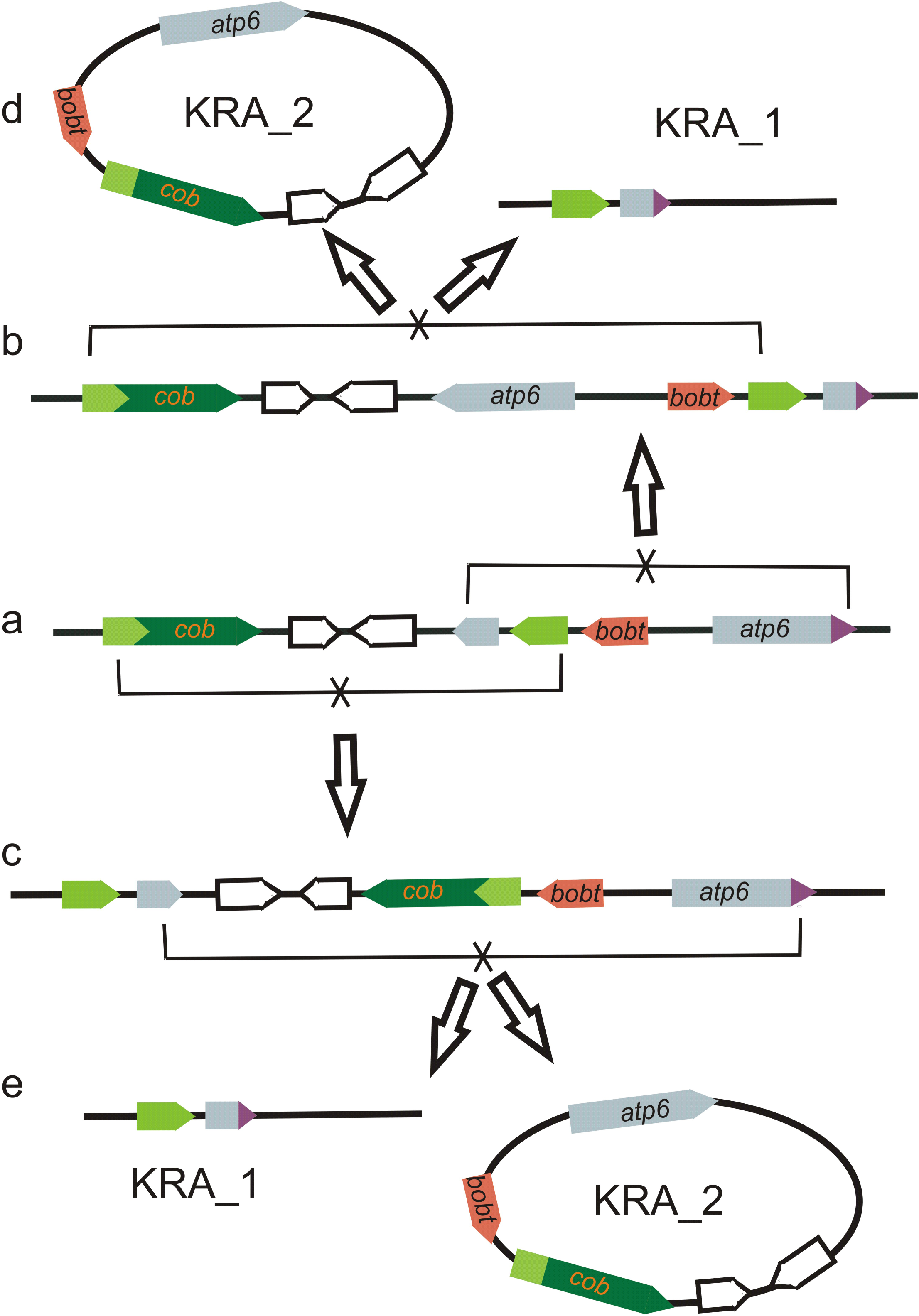
Diagram of homologous recombination across *cob* and *atp6*. The *cob* and partial *cob* sequences are green; the *atp6* and partial *atp6* sequences are grey. The conformation with all the genes located on the same DNA molecule (the joint chromosome KRA_1 and KRA_2) is depicted in the middle of the chart (a). *atp6* and partial *atp6* as well as *cob* and partial *cob* are in inverted orientation. Recombination across inverted repeats leads to the inversion between them; the DNA molecule is preserved. After the inversion, either the *cob* (top - b) or the *atp6* (bottom - c) repeats become directly oriented. The recombination across them generates an independent circular molecule, the semiautonomous chromosome KRA_2 (d, e). The *cob* gene is located downstream of the chimeric gene *bobt* on the chromosome KRA_2, and it is co-transcribed with it. In contrast, the *cob* gene is under the control of its own promoter in two conformations of the joint chromosome.

Homologous recombination across *atp6* or *cob* repeats not only maintains ‘recombinational equilibrium’ [47] between either separate or merged conformations of chromosome 1 and chromosome 2. It also profoundly impacts transcription of the *cob* gene. This essential gene is placed downstream of chimeric gene *bobt* and they are co-transcribed when collocated on chromosome 2, whereas it represents an independent transcription unit in two of the three possible recombinational configurations of the joint chromosome (Fig. 6a, b; Additional file 5: Figure S4 d). Thus, homologous recombination releases *cob* from the transcriptional control of *bobt*, a candidate CMS gene in *S*. *vulgaris* [46].

We performed Southern hybridizations to confirm the existence of various genomic configurations of the *cob* gene and to get approximate estimates of their abundances. The *cob* gene with its genuine promoter (similar to the *cob* promoter regions of the other *S. vulgaris* haplotypes) is present on the *Bgl*II fragment 6.57 kb, whereas the *bobt*-*cob* co-transcription unit is placed on the *Bgl*II fragment 5.73 kb. The *Bgl*II fragments 2.75 kb and 1.91 kb correspond to two variants containing a partial *cob* sequence (Fig. 7). Most plants showed higher intensity of the 5.73 kb band corresponding to the *bobt*-*cob* co-transcription unit. The intensities of the 6.57 kb and 5.73 kb bands varied across individuals, which suggests that the proportion of particular *cob* configuration is variable among individual plants and does not correlate with gender. The occurrence of recombination across *cob* and *atp6* repeats was also confirmed by Southern hybridization of genomic DNA digested with *Eco*RI (Additional file 7: Figure S5).

**Figure 7.**
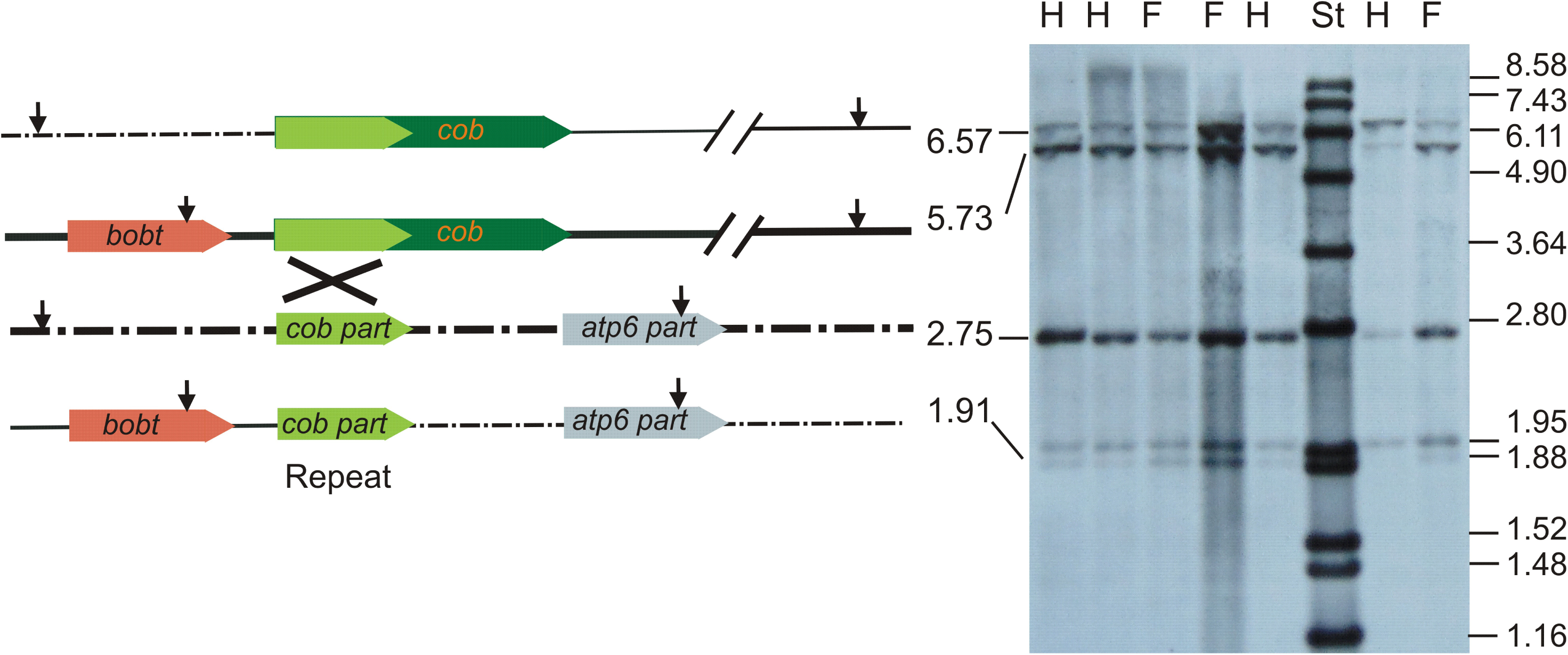
Recombination across *cob* repeat. The positions of the *Bgl*II sites in the vicinity of cob sequences are shown for four recombinant configurations. Southern blot probed with *cob* sequences is shown on the right. Total DNA was extracted from leaf tissues of three female and four hermaphroditic plants and digested with *Bgl*II. The sizes of the fragments corresponding to the respective recombinant configurations are given in kb. The sizes of the molecular standard in kb are shown on the right.

### Mitochondrial transcriptome of S. vulgaris KRA

The KRA mt genome contains five chimeric ORFs > 300 bp (Additional file 8: Figure S6), which represent CMS candidate genes. To evaluate their expression in the context of an entire mt genome in F and H individuals, we constructed a mt transcriptome of this haplotype. We compared depth of coverage and transcript editing in F and H plants, full sibs from the same cross, to reveal differentially expressed (DE) genes or genomic regions associated with CMS in the KRA haplotype.

We generated RNA-seq data from total RNA extracted from flower buds of three F and three H plants of *S. vulgaris* KRA. After rRNA removal and RNA fragmentation, all six samples were sequenced in a single HiSeq 4000 Illumina run, which produced approximately 38 M PE reads per sample; average fragment size was 192 nt. On average, 85% of read pairs passed quality filtering. About 5.2% of clean read pairs per sample were successfully mapped against the KRA mt genome.

### Differential gene expression between H and F individuals

We examined read coverage of mt protein genes, ORFs > 300 bp, and intergenic transcribed regions (‘transcription islands’). Because rRNA was eliminated before cDNA library construction and small RNAs (< 100 nt) were lost during RNA extraction, their DOC (depth of coverage) was not evaluated.

The *atp1* and *atp9* genes were the most highly expressed features, whereas *rpl5* and the plasmid-derived *dpo* gene showed little or no coverage, suggesting these genes may not be functional (Additional file 6: Data Set 1). Unlike other genes, *matR* was not covered evenly, a sharp drop in DOC was observed in the middle part of this gene as previously reported in KOV [45]. The marginal parts of *matR* encode the RT and X domains and they may exist as two separate mRNAs in mt transcriptomes of *S. vulgaris*. DOC of introns was on average lower than DOC of exons. However, some introns were covered to the same level as adjacent exons, e.g. *nad5* intron 4 (Additional file 5: Figure S4 b).

DOC was consistent across all six individuals and in general similar between F and H plants in the *S. vulgaris* haplotype KRA. The *bobt* gene harboring a 708 bp long chimeric ORF composed of portions of the *atp1* and *cox2* genes and of unknown sequence was highly DE (Additional file 8: Figure S6). This chimera was reported to be associated with CMS in the *S. vulgaris* haplotypes KRA and MTV [46]. We found > 3-fold higher DOC of the *bobt*-specific region in F than in H plants. The difference in the coverage between the genders was lower when an entire gene was considered owing to the *atp1*- and *cox2*-derived reads mapping to the homologous parts of the chimera (Additional file 6: Data Set 1). We validated abundances of *bobt* transcripts in 20 F and 20 H plants from three different controlled crosses by means of RT qPCR. The transcript levels were significantly higher in F than in H individuals, both in leaves and flower buds (ANOVA, p < 0.001), whereas the DNA copy number of the *bobt* gene was the same in the two genders and organs (Fig. 8).

**Figure 8.**
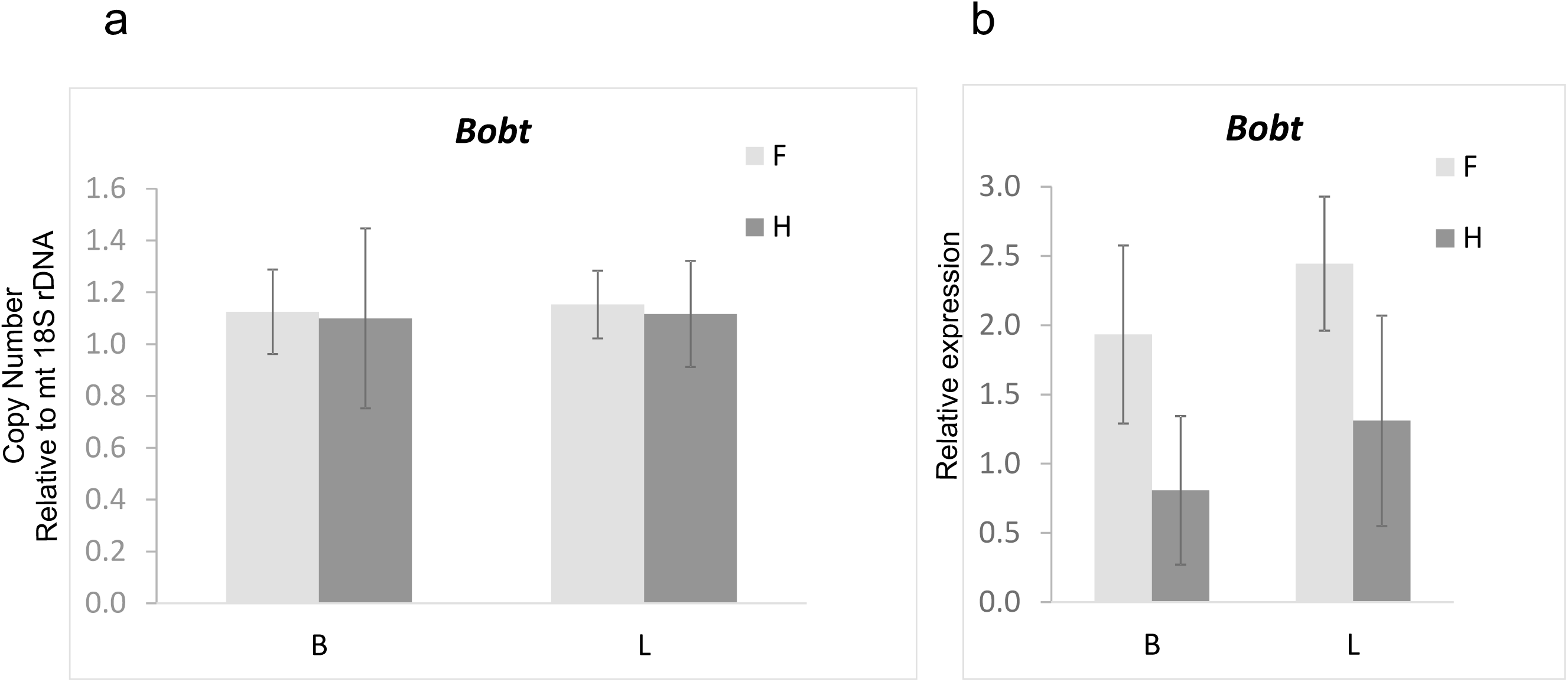
Copy numbers and expression of the chimeric gene *bobt*. Copy numbers were estimated with qPCR relative to mt 18S rDNA (a); relative expression was normalized with mt 18S rRNA (b).

The long chimeric ORF (1029 bp) contains the 5′ portion of the *cob* gene, which represents one of two repeat pairs enabling the recombination between chromosome 1 and chromosome 2. It is placed downstream of *bobt* in two of three possible configurations of the joint chromosome (Fig. 6). Because the DOC estimation is biased by *cob*-derived reads, only the portion of the sequence unique to this ORF was considered when analyzing this region. About 7-fold higher DOC was found in F than in H plants (Additional file 6: Data Set 2). The 1029-bp ORF is co-transcribed with *bobt* in some configurations, therefore a higher transcription from the *bobt* promoter is most likely responsible for the higher expression of this chimeric ORF in F than H plants. However, the coverage of this region was low even in F individuals (8-fold lower than *bobt* coverage), which makes possible function of this chimeric ORF in CMS less probable.

DOC was also very low for two other chimeric ORFs – 1.119388(orf)(230) and 1.231562(orf)(292) (Additional file 6: Data Set 2). The last of five chimeric ORFs under investigation - 247212(orf)(154) – was highly transcribed owing to its location just downstream of *cox1* and showed no differences in DOC between the genders. None of the additional 34 ORFs > 300 bp showed differential expression that would be consistent with a role as a CMS gene. Therefore, we can conclude that the *bobt* gene is the most likely CMS candidate among the ORFs under study in the *S. vulgaris* haplotype KRA.

There are 33 ‘transcription islands’ in the KRA genome. Some of them (173375(2500), 258075(300), or 287325(500)) exhibited as much as a 2-to 3-fold difference in DOC between F and H plants. Thus, it is possible that the RNAs encoded by these features could play a role in CMS. The highest difference in DOC between the genders among mt protein genes was recorded in *ccmFn*, which had about 3-fold more coverage in H than in F. This distinction is similar to the difference in *ccmFn* expression between F and H plants reported in *S. vulgaris* haplotype KOV [45].

### RNA editing differences between two mt genomes of S. vulgaris

We identified 417 unique C to U editing sites in the mt genome of *S. vulgaris* KRA, 302 of them located in protein-coding regions, 16 of them in type II introns. The remaining 99 edits were found in UTRs, ORFs, ‘transcription islands’, or in intergenic regions that were not classified as ‘transcription islands’. Editing sites in rRNAs and tRNAs were not evaluated due to their biased coverage influenced by rRNA elimination and tRNA loss in RNA extraction. Two hundred sixty-three editing sites in protein-coding genes were non-synonymous. Editing rates were similar between F and H plants (Additional file 6: Data Set 4).

We compared the comprehensive mt editome of *S. vulgaris* KRA with the previously published analysis of editing sites in the haplotype *S. vulgaris* KOV [47]. The loss of editing site caused by the substitution of C to T were recorded in the *nad5, ccmB* and *ccmFc* genes in KRA. The corresponding sites are edited in *S. latifolia*, whereas they have been lost in the haplotypes of *S. vulgaris* MTV, SD2, and S9L. All three sites are non-synonymous, but the loss of editing does not result in a change in protein sequence because of the C to T substitution in the DNA sequence. In addition, a loss of editing was found in the *atp6* portion of the chimeric ORF 1.119388(orf)(230). Interestingly, the editing site in the homologous position of the *atp6* gene was preserved. New editing was revealed in four sites. Their rate of editing was low and except for the edit in *ccmC*, they were silent, not affecting amino acid sequence (Table 3).

**Table 3.**
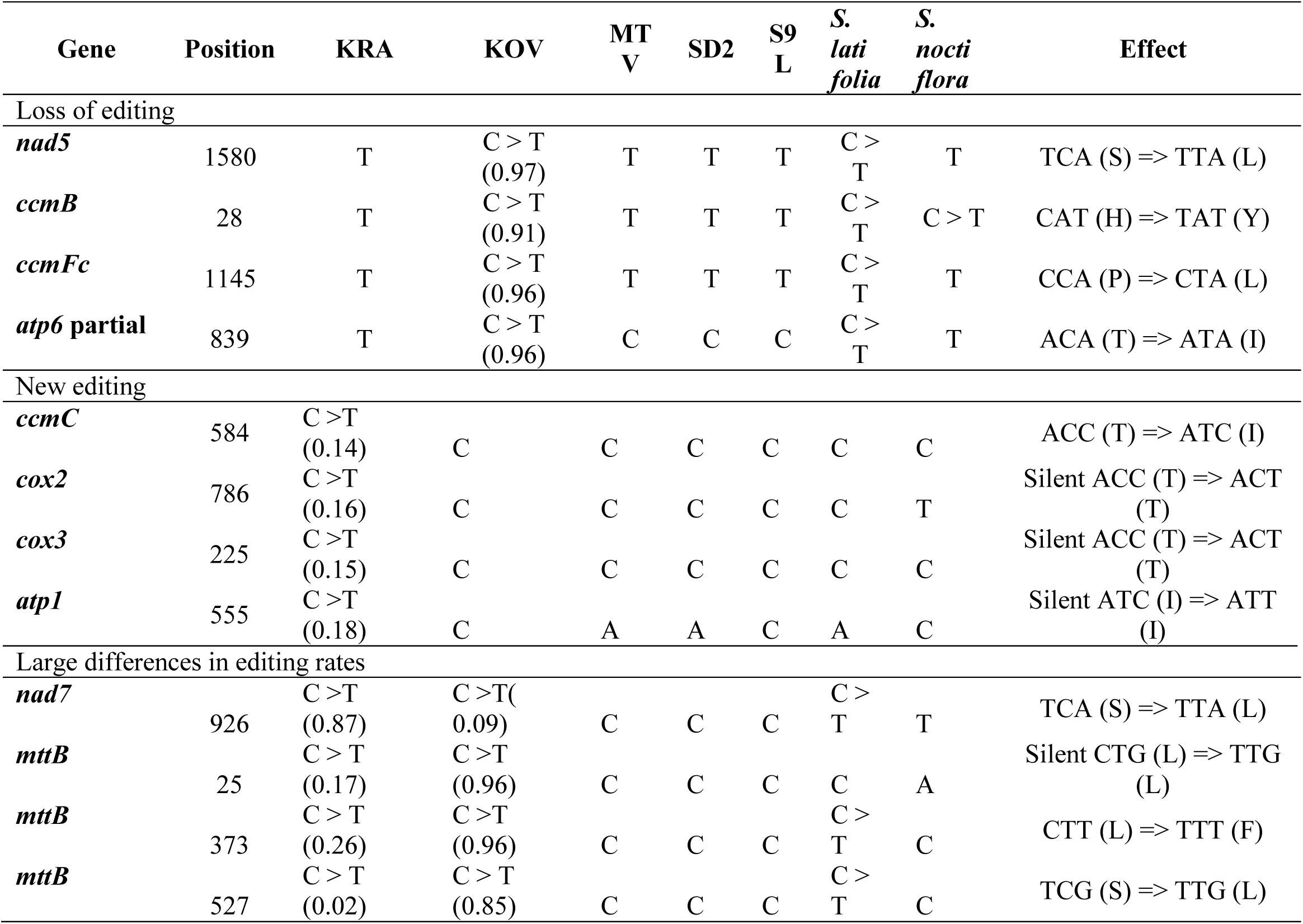
Differences in editing in protein coding genes between *S. vulgaris* mt genomes KOV and KRA. The sequence/editing in the corresponding positions in three additional *S. vulgaris* mt genomes and in two *Silene* species are shown. Editing in the MTV, SD2, and S9L haplotypes was not investigated; only sequence information is given. Editing rate, if available, is stated in parentheses. Editing rates higher than 0.8 in one haplotype and lower than 0.3 in the other one were considered highly different.

In general, there was a correlation in editing rates between *S. vulgaris* KRA and *S. vulgaris* KOV in both synonymous and non-synonymous sites (Additional file 9: Figure S7), with higher editing rates in non-synonymous sites, but several outliers were discovered. Three of four sites with highly distinct editing rates were non-synonymous, changing the amino acid composition of encoded proteins (Table 3). It means that, despite identical DNA sequence, the Nad7 and MttB proteins can differ between *S. vulgaris* KRA and *S. vulgaris* KOV. The *mttB* gene is the most variable gene between the two haplotypes in terms of editing, with three highly differentially edited positions.

The MttB protein interacts with a nucleus-encoded TatB subunit [48, 49]. The variation in amino acid sequence of the mt subunit may affect its interaction with the partner nuclear subunit. To determine whether within-species polymorphism exists in the nuclear *tatB* gene, we retrieved *tatB* sequences from cytoplasmic portions of the KRA and KOV transcriptomes of *S. vulgaris*. We identified 14 segregating sites among 12 individuals, 8 sites were non-synonymous. The number of synonymous SNPs per synonymous site was 0.035, the number of non-synonymous SNPs per non-synonymous site was 0.014, as estimated by DnaSP. Only a single allele was shared by the KOV and KRA plants, each individual was heterozygous. Polymorphism in the nuclear *tatB* gene may have compensated the variation in the MttB protein generated by editing of invariant primary transcripts of *mttB*.

Editing rates in intronic positions between the KRA and KOV haplotypes are highly correlated (Additional file 8: Figure S6 c). This supports the function of highly edited sites in stabilizing secondary RNA structure of introns. The KRA mt genome harbors one extra editing site in *nad4* intron 2.3 which was lost in the KOV haplotype (Additional file 6: Data Set 5), but in neither of the other *S. vulgaris* haplotypes, nor in *S. latifolia*. Information about editing is missing for the remaining haplotypes, but the high editing rate (0.84) in KRA and the substitution to a T in KOV suggests a functional importance of this position.

The KOV and KRA haplotypes share only about 50% of their overall sequence content and only one third of intergenic regions. Despite a high sequence divergence, they contain transcribed intergenic regions with homologous editing positions. Interestingly, a moderately edited position in the ‘transcription island’ 1.371387(450) in the KRA haplotype was lost in the KOV haplotype (Table 4). Because this C to T substitution is the only polymorphism in 200 nt long homologous region, editing in this position in the KRA genome may not be a side effect of editing machinery operating primarily at other sites, but it may be necessary to fulfill some function. Thus, the C to T substitution could reflect selection to conserve this function, or it could have been mediated by retroprocessing of the edited transcript, which is replacement of genomic sequence with cDNA generated by reverse transcription of the edited transcript [50, 51].

## Discussion

### Small chromosomes of the mt genome KRA resemble circular bacterial plasmids

The mt genome of the Asian accession of *S. vulgaris* KRA collected near Krasnoyarsk (Russia) [46] represents the fifth completely sequenced mt genome of this gynodioecious species. It further expands an extraordinarily large range of mtDNA variation in *S. vulgaris*. About 14% of the KRA mt genome is unique, not matching any GenBank record, including the other four mt genome sequences from this species. It is the most similar to the S9L mt genome originating from plants collected in Virginia (USA), and least similar to the KOV mt genome from the European population growing near Prague (Czech Republic) [9]. The close relationship between the KRA and S9L mt genomes can be explained by human transport [52], which introduced the ancestor of *S. vulgaris* S9L from a site in Eurasia to Virginia on the opposite side of globe.

The multichromosomal mt genomes of KRA and S9L share one autonomous chromosome 9.5 kb in length (chromosome 6 in S9L, chromosome 3 in KRA) which is nearly identical in sequence (99.94%) between the two mt genomes, differing only by 6 SNPs. This high level of similarity contrasts with the second homologous pair – small chromosome 7 in S9L (JQ771316, 6.5kb) and chromosome 4 in KRA (MH455605, 2.6 kb). They differ by large indels and their homologous regions are similar only between 94% and 97% nucleotide identity. Supposing that homologous chromosomes originate from the common ancestor of KRA and S9L and, thus, have both been diverging for the same amount of time, the discrepancy in evolutionary rate between the two pairs of homologous chromosomes is enormous. Evolutionary rates are known to vary among mt protein coding genes [53], including *atp9* evolving faster than other mt genes in *Silene* [26]. The rate of sequence evolution was also found to be very high in mitochondrial plasmids [54], which may be explained if they use a distinct replication/repair machinery. Accordingly, small chromosomes may be replicated/repaired differently from the rest of mt genome including the autonomous 9.5 kb chromosome. Whereas chromosome 3 in KRA (9.5 kb) shows an equimolar ratio with the main chromosome, the copy number of small chromosome 4 in KRA was about 4-to 8-fold higher, with more in leaves that in flower buds. The differences in the molar ratio of small and large mt chromosomes between the plant organs could be another signature of a distinct mode of DNA replication or maintenance. However, unlike mitochondrial plasmids, neither chromosome 4 in KRA nor its homologs in the other sequenced mt genomes of *S. vulgaris* encode any proteins. Thus, they likely rely on nuclear-encoded replication and repair machinery like the rest of the mt genome.

Not only the copy numbers, but also the structure of chromosome 4 differed between flower buds and leaves. Fragments corresponding to supercoiled DNA prevailed in buds, whereas relaxed circles originating from supercoils by single-strand breaks similar to the structures produced by nicking enzyme were dominant in leaves. The observed variation may be caused by differential damage between leaves and buds during DNA extraction, but they may also reflect differences in mtDNA structure between the two organs. The changes of physical structure of small chromosomes are in agreement with a general view of the modification of mtDNA integrity and amount during individual development reported by [55, 56].

Although the lack of coding capacity and any sequence similarity in general make it difficult to speculate about the role of small chromosomes in *S. vulgaris*, we may hypothesize that (1) they are selfish DNA elements utilizing replication machinery of plant mitochondria for their propagation, or (2) that they perform some function. The presence of small chromosomes in all five completely sequenced mt genomes of *S. vulgaris* from three continents and the conservancy of the occasionally transcribed areas seem to prefer the second possibility, although the maintenance of the small mt chromosomes as a by-product of replication or recombination cannot be ruled out.

Unlike other sequenced mt genomes of *S. vulgaris*, the KRA mt genome contains another small chromosome, which is only 1.5 kb long. Its sequence is unique, providing no match with any GenBank record including other *S. vulgaris* mt sequences. Despite sharing a 280-bp repeat with main chromosome 1, the small chromosome 5 appears to be autonomous. No evidence for recombination between it and the main chromosome was found, neither in RNA-seq nor Southern hybridization data. The structure of chromosome 5 is similar to chromosome 4 and to bacterial plasmids, but unlike multimeric chromosome 4, only the monomers of chromosome 5 were observed. Because chromosome 5 is present only in the KRA mt genome, it is dispensable in other mt genomes and may be less likely to code for any function.

Small autonomous mt chromosomes of *S. vulgaris* differ from plasmids described in plant mitochondria including *S. vulgaris* [57, 58]. They contain no coding sequences, and do not recombine with other mt chromosomes. Their structure resembles bacterial plasmids occurring in the form of supercoiled, linear or relaxed circular DNA. The copy number of small autonomous mt chromosomes is more variable and higher than copy number of large mt chromosomes. Small autonomous mt chromosomes of *S. vulgaris* therefore represent a specific form of plant mtDNA with so far unknown origins and function.

### Recombination in the KRA mt genome

The 708 bp long chimeric ORF identified as the CMS candidate gene *bobt* in *S. vulgaris* [46] was a highly DE feature between F and H plants in the KRA mt transcriptome. Four other chimeric ORFs were either expressed at very low levels or exhibited equivalent levels of transcript abundance in both genders, likely owing to a close proximity to a protein coding gene. These findings strengthen the hypothesized role of *bobt* as a CMS gene.

The *bobt* gene is located on chromosome 2 upstream of the *cob* gene with which it is co-transcribed. The co-transcription of CMS genes with essential protein-coding genes has been observed in several species. For example, orf256 is co-transcribed with *cox1* in wheat [59], and orf456 is co-transcribed with *cox2* in chili pepper [60]. The co-transcription of a CMS factor with an essential gene may constrain transcription suppression as a mechanism for fertility restoration because of the need to maintain sufficient production of the essential protein. Homologous recombination across the *cob* repeat in the KRA mt genome overcomes this constraint and releases the *cob* gene from co-transcription with *bobt*. The proportion of particular recombinational configurations of the *bobt* and *cob* genes varies among the plants but does not correlate with gender. It will be interesting to investigate, whether the ratio between the alternative variants can be modified by stress conditions or differs across developmental stages.

Plant mt genomes are highly dynamic, undergoing frequent intramolecular recombination across numerous repeats [10, 61], which may originate in the course of a double strand break repair [14, 18, 62]. Our results suggest that rearrangements of plant mt genomes mediated by homologous recombination may also influence transcriptional context of essential genes. A similar observation affecting the *cox2* gene in maize was published by [63]. Homologous recombination between a linear plasmid and mt chromosome, which led to the expression of *orf355/orf77* and male sterility was observed by [64]. Thus, recombination in plant mt genomes, which is often associated with DNA repair and replication, may also control the expression of vital mt genes.

### The comparison of the mt editomes in S. vulgaris

A comprehensive comparison of the mt editing sites between the mt haplotypes KRA and KOV revealed another layer of nucleotide variation in the mitochondria of *S. vulgaris*. The loss of editing sites observed in the genus *Silene* [65] has continued at the within-species level in *S. vulgaris*. The replacement of C for T in KRA did not change amino-acid sequence of encoded protein. However, it may affect the evolution of nuclear editing factors responsible for the recognition of the respective editing sites. If editing is not required anymore owing to C to T replacement, the corresponding editing factor may not experience selection and it may start to accumulate mutations. When editing sites are not conserved in the species, crossing may combine mt genome requiring editing at specific positions with nuclear backgrounds that contain editing factors that are less efficient at editing those sites. An example is the defect in plastid RNA editing in the nightshade/tobacco cybrid, which resulted in pigment deficiency [66].

Thus, within-species polymorphism in editing sites represents an example of nuclear-cytoplasmic interaction which may lead to incompatibilities and subsequently to reproduction barriers among the populations of the same species [67, 68, 69]. We may expect higher levels of polymorphism in nuclear editing factors in species with polymorphism in editing sites like *S. vulgaris*. We may also expect the existence of homologous editing sites with highly different editing rates among the haplotypes of this species. We found sites edited to highly different extents in KRA and KOV in the *mttB* and *nad7* genes despite the identical nucleotide sequences of the two genes between the KOV and KRA haplotypes. Three of the four highly differentially edited sites were non-synonymous. Two highly differentially edited non-synonymous sites were located in the *mttB* (also known as *tatC*) gene encoding the subunit of protein translocation complex located in inner mt membrane. We detected a high polymorphism in the nuclear *tatB* gene coding for the subunit interacting with MttB. Fast evolution of organellar genomes may select for compensatory mutations in interacting proteins encoded by nucleus [70, 71, 72]. For example, nucleus-encoded subunits of organellar ribosomes have higher amino acid sequence polymorphism than their cytosolic counterparts in species with rapid mt and plastid genome evolution [73]. The *mttB* genes are identical between KOV and KRA, the variation in amino acid sequences of their products was introduced by editing. Organellar editing increases the variation in mt proteins and may contribute to relaxed functional constraints of nucleus-encoded subunits. It should, therefore, be considered as an important factor in the co-evolution between nuclear and organellar genomes.

We have also observed sites with low editing extent, which did not pass the threshold values for editing in KOV [45]. They most often resulted in silent substitutions. The ongoing changes in editing sites in *S. vulgaris* follows the general pattern documented at much higher phylogenetic levels of angiosperms [74].

‘Transcription islands’ code for RNAs with zero or limited protein-coding capacity. Long non-coding RNAs > 150 nt were reported in mitochondria of *Arabidopsis* [75] and tobacco [76]. There is no information about the specific function of non-coding RNA encoded by plant mtDNA [77]. However, long non-coding RNA associated with CMS has been recently revealed in the haplotype KOV of *S. vulgaris* [45]. The existence of shared ‘transcription islands’ between the KRA and KOV mt transcriptomes and their conserved editing sites suggest possible functional importance of these regions encoding non-coding RNAs.

## Conclusions

Multichromosomal mt genomes of *S. vulgaris* exhibit an unprecedented level of intraspecific variation in sequence content. Inter-genomic recombination also played a major role in their structural and sequence evolution. However not all chromosomes participate in the recombining genetic pool. Small circular non-recombining gene-less chromosomes resembling bacterial plasmids in structure are present in all five completely sequenced mt genomes of *S. vulgaris*, which suggests their possible, albeit unknown function.

We report the formation of a co-transcriptional unit by homologous recombination, placing the gene *cob* either under or outside the control of the promoter of the chimeric gene *bobt* in the KRA haplotype. In this manner, the proportion of *cob* copies transcribed from the promoter of the CMS candidate gene *bobt* is influenced by recombination. This observation illustrates the role of homologous recombination in the control of mt gene transcription, and possibly also in CMS.

Genetic diversity of mt genomes of *S. vulgaris* is accompanied by the diversity in mt editing sites. The independent losses of three editing sites and the existence of positions with highly different editing rates were detected by the comparison of mt transcriptomes of two haplotypes of *S. vulgaris*. These findings bring evidence about fast evolution of editing sites even within a single species.

## Methods

### Plant material

Seeds of *Silene vulgaris* KRA haplotype were collected in a site near Krasnoyarsk (Siberia, Russia) in 2010 [46]. A single F plant was pollinated by an H plant of the same population. The progeny were cultivated in the Institute of Experimental Botany (IEB) greenhouse under supplemental lighting (16/8 h light/ dark) in pots filled with perlite, vermiculite, and coconut coir (1:1:1), and fertilized 2–3 times per week. Three F and three H full-sib individuals were selected and tested for homoplasmy by amplifying, cloning, and sequencing the highly polymorphic *atp1* gene, using the pGEM T-easy vector (Promega, WI, USA). The sequences of all 50 clones from each individual exhibited no differences.

### Mt genome assembly and sequence analysis

The mtDNA extraction procedure was described previously [9]. Briefly, about 2 g of flower buds (1-3 mm) were ground in grinding buffer and the suspension was filtered through Miracloth and centrifuged. The supernatant was centrifuged at higher speed (12 000g), and the pellet was immediately utilized to isolate DNA. MtDNA was used to generate 3kb paired-end library and sequenced on one-half of a plate on a Roche 454 GS_FLX platform with Titanium chemistry at the DNA Sequencing Center at Brigham Young University. The genome assembly and annotation generally followed the procedure described by [2] using Roche’s GS de novo Assembler v2.6 (‘Newbler’). After performing an initial assembly, the reads mapping to contigs with > 80x coverage were retrieved and reassembled. The KRA mt genome was annotated according to previously published mt genomes of *S. vulgaris* [9], tRNA genes were searched using tRNAscan [78]. The annotated genome sequence was deposited in GenBank under the numbers MH455602-MH455606, and visualized by the OGDRAW software tools [79]. The sizes and repeats of the KRA chromosomes were visualized by Circos v.0.69 [80]. Repetitive sequences, plastid-derived regions and shared sequence content were estimated as described by [2]. ORFs > 300 bp were detected in Geneious 7.1.5. Phylogenetic relationships were inferred using multiple nucleotide sequence alignment of concatenated protein-coding genes extracted from completely sequenced mt genomes of *S. vulgaris* [9, 81] and mt genomic draft of *S. vulgaris* subsp. *prostrata* D11 (GenBank accession MH576576) generated by MUSCLE [82] implemented in Geneious 7.1.5. The alignment was analyzed by the maximum-likelihood (ML) method using RAxML [83]. A gamma distribution of rate heterogeneity was applied, and bootstrap support of the ML tree was calculated from 1000 pseudoreplicates. The numbers of synonymous and non-synonymous SNPs in the *tatB* gene were estimated by DnaSP v5 [84].

### Southern blot hybridizations

We performed Southern hybridization to analyze genome structure of small autonomous chromosomes and to demonstrate recombination events across the *cob* and *atp6* genes. Total genomic DNA was isolated by a sorbitol extraction method [85] from leaves or flower buds flash-frozen in liquid nitrogen. Samples containing about 8 μg of DNA were digested with either a restriction endonuclease (*Bgl*II-HF or *Eco*RI-HF) or with a single-strand cleaving nickase (Nb.*Bss*SI [chromosome 4 specific] or Nb.*Bsr*DI [chromosome 5 specific]) (New England BioLabs, Frankfurt, Germany). Additional samples were left undigested as a control. The samples were electrophoresed overnight on a 0.9% agarose gel and capillary blotting was performed as described previously [9]. Probe targeting the regions on chromosomes 4 and 5 or on the *cob* gene were PCR amplified from genomic DNA (Additional file 6: Data Set 6), labeled with digoxigenin (DIG), and hybridized as described previously [81].

### RNA extraction and Illumina sequencing

Strand-specific cDNA libraries were prepared from total RNA after rRNA depletion using the Epicentre Ribo Zero Plant Leaf kit (Cat No. RZPL1224) according to the manufacturer’s protocol. All six specimens were sequenced at University of California Davis Genome Center (USA) on a single lane Illumina HiSeq 4000, which generated paired-end reads (2 x 150 cycles). The reads were trimmed using Trimmomatic 0.32 [86] in paired-end mode with a quality threshold of 20. Reads with a post-trim length less than 140 were removed. Approximately 5.3% read pairs of the entire data set were removed by trimming.

### Transcriptome analysis

Initial alignment was performed using GSNAP v. 2014-12-23 [87] in paired-end mode using known splice junctions [45] as *a priori* information for GSNAP. Read mapping, variant site discovery by HaplotypeCaller module in GATK 3.4 [88] and transcript abundance estimation followed the procedures described in detail by [45]. Coverages were normalized as Transcripts Per Kilobase Million (TPM) [89]. Transcribed sequences in intergenic regions (‘transcription islands’) were identified using the makewindows function in bedtools v 2.24.0 [90] to generate 100-bp sliding windows across the mt KRA genome and to compare intergenic and coding regions [45]. The threshold for ‘transcription islands’ was approximately 12000 reads per window in the merged bam file for all samples, which was equivalent to between 1700-2200 reads DOC depending on the sample.

### Transcript abundance and gene copy number estimation by RT qPCR and qPCR

Complementary DNA (cDNA) was synthesized using Transcriptor HF Reverse Transcriptase (Roche Applied Science, Mannheim, Germany) and random hexamers as described by [45]. Three independent RT reactions were set up for each RNA sample. Quantitative PCR was performed using Light Cycler 480 SYBR Green I Master on a LightCycler 480 instrument (Roche Applied Science, Mannheim, Germany). Reaction mixture contained 5 µl 2× MasterMix, primers in specific concentrations (Additional file 6: Data Set 6) and 2.5 µl of 20× diluted first-strand cDNA in a total volume of 10 µl. Cycling conditions were the same as described by[45]. Each cDNA sample was measured three times, and means and standard deviations were calculated from six values (2 cDNAs × 3 measurements). Gene copy number was estimated by qPCR in the same way as transcript abundance. The intragenomic ratio of mt gene copy numbers was estimated relative to the mt *rrn18* gene. The ratio between mt DNA and nuclear DNA was measured relative to nuclear 18S rDNA. Each estimation was repeated four times.

## Additional files

**Additional file 1. Figure S1.** Maps of *S. vulgaris* KRA mitochondrial chromosomes.

**Additional file 2. Figure S2.** Phylogenetic relationships among five mitochondrial genomes of *S. vulgaris*.

**Additional file 3. Table S1.** Nucleotide polymorphism in the *atp6* gene.

**Additional file 4. Figure S3**. The alignment of the small mitochondrial autonomous chromosomes from five genomes of *S. vulgaris.*

**Additional file 5. Figure S4**. Coverage of selected mitochondrial genes of *S. vulgaris* KRA.

**Additional file 6**. **Data Set S1**. Coverage Genes. **Data Set S2.** Coverage ORFs. **Data Set S3.** Coverage Intergenic. **Data Set S4**. Editing Sites**. Data Set S5**. Editing Comparison. **Data Set S6.** Primers.

**Additional file 7. Figure S5**. Recombination across the *cob* repeat visualized by Southern hybridization with *Eco*RI digested DNA.

**Additional file 8. Figure S6.** Summary of chimeric ORFs > 300 bp in length in the mitochondrial genome of *S. vulgaris* KRA.

**Additional file 9**. **Figure S7.** Comparison of editing extent between *S. vulgaris* KRA and KOV.

## Acknowledgements

The authors thank Pavla Koloušková for excellent technical assistance in the lab and in the greenhouse. The bioinformatic help provided by Miloslav Juříček and Manuela Krüger is highly appreciated, as well as the assitance of Filip Štorch with graphical design of some figures. Access to computing and storage facilities owned by parties and projects contributing to the National Grid Infrastructure MetaCentrum, provided under the programme “Projects of Large Infrastructure for Research, Development, and Innovations” (LM2010005), is greatly appreciated.

## Availability of data and materials

The data of this study data have been deposited in the NCBI with BioProject accession number PRJNA321915. The reads from hermaphrodites of *S. vulgaris* KRA can be found under the number SRS2438489, the reads from females under the number SRS2438490. Mitochondrial genome assembly of *S. vulgaris* KRA has been deposited to GenBank with accession numbers MH455602-MH455606. Mitochondrial genomic draft of *S. vulgaris* subsp. *prostrata* D11 is stored in GenBank with the number MH576576.

## Authors’ contributions

HŠ conceived and designed the experiments and made some analyzes. JDS performed most data analyzes, DBS designed data analyzes and provided trouble shooting, OA and KM performed the experiments, JW and MP participated in data analyzes. HŠ, DBS and JDS wrote the manuscript. All authors read and approved the final manuscript.

## Ethics approval and consent to participate

Not applicable

## Consent for publication

Not applicable

## References

1. Skippington E, Barkman TJ, Rice DW, Palmer JD. Miniaturized mitogenome of the parasitic plant Viscum scurruloideum is extremely divergent and dynamic and has lost all nad genes. Proc Natl Acad Sci U S A. 2015;112:E3515-24. doi:10.1073/pnas.1504491112.

2. Sloan DB, Alverson AJ, Chuckalovcak JP, Wu M, McCauley DE, Palmer JD, et al. Rapid evolution of enormous, multichromosomal genomes in flowering plant mitochondria with exceptionally high mutation rates. PLoS Biol. 2012;10.

3. Mower JP, Case AL, Floro ER, Willis JH. Evidence against equimolarity of large repeat arrangements and a predominant master circle structure of the mitochondrial genome from a monkeyflower (Mimulus guttatus) lineage with cryptic CMS. Genome Biol Evol. 2012;4:670–86.

4. Adams KL, Qiu Y-L, Stoutemyer M, Palmer JD. Punctuated evolution of mitochondrial gene content: High and variable rates of mitochondrial gene loss and transfer to the nucleus during angiosperm evolution. Proc Natl Acad Sci U S A. 2002;99:9905–12. doi:10.1073/pnas.042694899.

5. Richardson AO, Rice DW, Young GJ, Alverson AJ, Palmer JD. The “fossilized” mitochondrial genome of Liriodendron tulipifera: Ancestral gene content and order, ancestral editing sites, and extraordinarily low mutation rate. BMC Biol. 2013;11:29.

6. Allen JO, Fauron CM, Minx P, Roark L, Oddiraju S, Guan NL, et al. Comparisons among two fertile and three male-sterile mitochondrial genomes of maize. Genetics. 2007;177:1173–92.

7. Davila JI, Arrieta-Montiel MP, Wamboldt Y, Cao J, Hagmann J, Shedge V, et al. Double-strand break repair processes drive evolution of the mitochondrial genome in Arabidopsis. BMC Biol. 2011;9:64.

8. Darracq A, Varré JS, Maréchal-Drouard L, Courseaux A, Castric V, Saumitou-Laprade P, et al. Structural and content diversity of mitochondrial genome in beet: A comparative genomic analysis. Genome Biol Evol. 2011;3:723–36.

9. Sloan DB, Müller K, McCauley DE, Taylor DR, Storchová H. Intraspecific variation in mitochondrial genome sequence, structure, and gene content in Silene vulgaris, an angiosperm with pervasive cytoplasmic male sterility. New Phytol. 2012;196:1228–39. doi:10.1111/j.1469-8137.2012.04340.x.

10. Maréchal A, Brisson N. Recombination and the maintenance of plant organelle genome stability. New Phytol. 2010;186:299–317.

11. Small ID, Isaac PG, Leaver CJ. Stoichiometric differences in DNA molecules containing the *atpA* gene suggest mechanisms for the generation of mitochondrial genome diversity in maize. EMBO J. 1987;6:865–69.

12. Mackenzie SA, Chase CD. Fertility Restoration Is Associated with Loss of a Portion of the Mitochondrial Genome in Cytoplasmic Male-Sterile Common Bean. Plant Cell. 1990;2:905–12. doi:10.1105/tpc.2.9.905.

13. Chen J, Guan R, Chang S, Du T, Zhang H, Xing H. Substoichiometrically different mitotypes coexist in mitochondrial genomes of Brassica napus L. PLoS One. 2011;6: e17662.

14. Gualberto JM, Newton KJ. Plant Mitochondrial Genomes: Dynamics and Mechanisms of Mutation. Annu Rev Plant Biol. 2017;68:225–52. doi:10.1146/annurev-arplant-043015-112232.

15. Abdelnoor R V., Yule R, Elo A, Christensen AC, Meyer-Gauen G, Mackenzie SA. Substoichiometric shifting in the plant mitochondrial genome is influenced by a gene homologous to MutS. Proc Natl Acad Sci U S A. 2003;100:5968–73. doi:10.1073/pnas.1037651100.

16. Shedge V, Arrieta-Montiel M, Christensen AC, Mackenzie SA. Plant Mitochondrial Recombination Surveillance Requires Unusual RecA and MutS Homologs. Plant Cell. 2007;19:1251–64. doi:10.1105/tpc.106.048355.

17. Kuhn K, Gualberto JM. Recombination in the Stability, Repair and Evolution of the Mitochondrial Genome. In: Marechal Drouard L, editor. Mitochondrial Genome Evolution. Adv Bot Res. 2012;63:215–52.

18. Christensen AC. Genes and junk in plant mitochondria-repair mechanisms and selection. Genome Biol Evol. 2014;6:1448–53.

19. Suzuki N, Koussevitzky S, Mittler R, Miller G. ROS and redox signalling in the response of plants to abiotic stress. Plant Cell Environ. 2012;35:259–70.

20. Anand RP, Lovett ST, Haber JE. Break-induced DNA replication. Cold Spring Harb Perspect Biol. 2013;5:a010397.

21. Saini N, Ramakrishnan S, Elango R, Ayyar S, Zhang Y. Migrating bubble during break-induced replication drives conservative DNA synthesis. Nature. 2013;502:389–92.

22. Wolfe KH, Li W-H, Sharp PM. Rates of Nucleotide Substitution Vary Greatly among Plant Mitochondrial, Chloroplast, and Nuclear DNAs. Proc Natl Acad Sci U S A. 1987;84:9054–8. doi:10.1073/pnas.84.24.9054.

23. Palmer JD, Herbon LA. Plant Mitochondrial DNA Evolves Rapidly in Structure, but Slowly in Sequence. J Mol Evol. 1988;28:87–97.

24. Cho Y, Mower JP, Qiu Y-L, Palmer JD. Mitochondrial substitution rates are extraordinarily elevated and variable in a genus of flowering plants. Proc Natl Acad Sci U S A. 2004;101:17741–6. doi:10.1073/pnas.0408302101.

25. Mower JP, Touzet P, Gummow JS, Delph LF, Palmer JD. Extensive variation in synonymous substitution rates in mitochondrial genes of seed plants. BMC Evol Biol. 2007;7:7.

26. Sloan DB, Oxelman B, Rautenberg A, Taylor DR. Phylogenetic analysis of mitochondrial substitution rate variation in the angiosperm tribe Sileneae. BMC Evol Biol. 2009;9:260.

27. Hanson M, Bentolila S. Interactions of mitochondrial and nuclear genes that affect male gametophyte development. Plant Cell. 2004;16:154–70. doi:10.1105/tpc.015966.

28. Horn R, Gupta KJ, Colombo N. Mitochondrion role in molecular basis of cytoplasmic male sterility. Mitochondrion. 2014;19 PB:198–205. doi:10.1016/j.mito.2014.04.004.

29. Touzet P, Meyer EH. Cytoplasmic male sterility and mitochondrial metabolism in plants. Mitochondrion. 2014;19 PB:166–71. doi:10.1016/j.mito.2014.04.009.

30. Hu J, Huang W, Huang Q, Qin X, Yu C, Wang L, et al. Mitochondria and cytoplasmic male sterility in plants. Mitochondrion. 2014;19 PB:282–8. doi:10.1016/j.mito.2014.02.008.

31. McCauley DE, Olson MS. Do recent findings in plant mitochondrial molecular and population genetics have implications for the study of gynodioecy and cytonuclear conflict? Evolution. 2008;62:1013–25.

32. Guyon, P.H; Vichot P; VanDamme J. Nuclear-Cytoplasmic Male Sterility?: Single-Point Equilibria Versus Limit Cycles. Am Nat. 1991;137:498–514.

33. Dufaÿ M, Touzet P, Maurice S, Cuguen J. Modelling the maintenance of male-fertile cytoplasm in a gynodioecious population. Heredity. 2007;99:349–56.

34. Charlesworth D. A further study of the problem of the maintenance of females in gynodioecious species. Heredity. 1981;46:27–39.

35. Städler T, Delph LF. Ancient mitochondrial haplotypes and evidence for intragenic recombination in a gynodioecious plant. Proc Natl Acad Sci U S A. 2002;99:11730–35.

36. Touzet P, Delph LF. The effect of breeding system on polymorphism in mitochondrial genes of silene. Genetics. 2009;181:631–44.

37. Delph LF, Montgomery BR. The Evolutionary Dynamics of Gynodioecy in *Lobelia*. Int J Plant Sci U S A 2014;175:383–91. doi:10.1086/675572.

38. Lahiani E, Dufaÿ M, Castric V, Le Cadre S, Charlesworth D, Van Rossum F, et al. Disentangling the effects of mating systems and mutation rates on cytoplamic diversity in gynodioecious Silene nutans and dioecious Silene otites. Heredity. 2013;111:157–64.

39. Štorchová H, Olson MS. Comparison between mitochondrial and chloroplast DNA variation in the native range of Silene vulgaris. Mol Ecol. 2004;13:2909–19.

40. Houliston GJ, Olson MS. Nonneutral evolution of organelle genes in Silene vulgaris. Genetics. 2006;174:1983–1994.

41. Barr CM, Keller SR, Ingvarsson PK, Sloan DB, Taylor DR. Variation in mutation rate and polymorphism among mitochondrial genes of Silene vulgaris. Mol Biol Evol. 2007;24:1783– 1791.

42. Ingvarsson PK, Taylor DR. Genealogical evidence for epidemics of selfish genes. Proc Natl Acad Sci U S A 2002;99:11265–11269. doi:10.1073/pnas.172318099.

43. Olson MS, McCauley DE. Mitochondrial DNA diversity, population structure, and gender association in the gynodioecious plant *Silene vulgaris*. Evolution. 2002;56:253–62.

44. Sanderson BJ, Augat ME, Taylor DR, Brodie ED. Scale dependence of sex ratio in wild plant populations: Implications for social selection. Ecol Evol. 2016;6:1411–19.

45. Stone JD, Koloušková P, Sloan DB, Štorchová H. Non-coding RNA may be associated with cytoplasmic male sterility in Silene vulgaris. J Exp Bot. 2017;68:1599–1612.

46. Štorchová H, Müller K, Lau S, Olson MS. Mosaic origins of a complex chimeric mitochondrial gene in Silene vulgaris. PLoS One. 2012;7:e30401

47. Lonsdale DM, Brears T, Hodge TP, Melville SE, Rottmann WH. The plant mitochondrial genome - homologous recombination as a mechanism for generating heterogeneity. Philos Trans R Soc B-Biol Sci. 1988;319:149–63.

48. Carrie C, Weißenberger S, Soll J. Plant mitochondria contain the protein translocase subunits TatB and TatC. J Cell Sci. 2016;129:3935–47. doi:10.1242/jcs.190975.

49. Arenas A, Gonzalez-Duran E, Gomez I, Burger M, Brennicke A, Takenaka M et al. The Pentatricopeptide Repeat Protein MEF31 is Required for Editing at Site 581 of the Mitochondrial tatC Transcript and Indirectly Influences Editing at Site 586 of the Same Transcript. Plant Cell Physiol. 2018;59:355–65.

50. Cuenca A, Petersen G, Seberg O, Davis JI, Stevenson DW. Are substitution rates and RNA editing correlated? BMC Evol Biol. 2010;10:349. doi:10.1186/1471-2148-10-349.

51. Hecht J, Grewe F, Knoop V. Extreme RNA editing in coding islands and abundant microsatellites in repeat sequences of Selaginella moellendorffii mitochondria: The root of frequent plant mtDNA recombination in early tracheophytes. Genome Biol Evol. 2011;3:344–58.

52. Keller SR, Fields PD, Berardi AE, Taylor DR. Recent admixture generates heterozygosity-fitness correlations during the range expansion of an invading species. J Evol Biol. 2014;27:616– 27.

53. Zhu A, Guo W, Jain K, Mower JP. Unprecedented heterogeneity in the synonymous substitution rate within a plant genome. Mol Biol Evol. 2014;31:1228–36.

54. Warren JM, Simmons MP, Wu Z, Sloan DB. Linear plasmids and the rate of sequence evolution in plant mitochondrial genomes. Genome Biol Evol. 2016;8:364–74.

55. Oldenburg DJ, Kumar RA, Bendich AJ. The amount and integrity of mtDNA in maize decline with development. Planta. 2013;237:603–17.

56. Oldenburg DJ, Bendich AJ. DNA maintenance in plastids and mitochondria of plants. Front Plant Sci. 2015;6:883. doi:10.3389/fpls.2015.00883.

57. Andersson-Ceplitis H, Bengtsson BO. Transmission rates and phenotypic effects of mitochondrial plasmids and cytotypes in Silene vulgaris. Evolution. 2002;56:1586–91

58. Handa H. Linear plasmids in plant mitochondria: Peaceful coexistences or malicious invasions? Mitochondrion. 2008;8:15–25.

59. Hedgcoth C, El-Shehawi AM, Wei P, Clarkson M, Tamalis D. A chimeric open reading frame associated with cytoplasmic male sterility in alloplasmic wheat with Triticum timopheevi mitochondria is present in several Triticum and Aegilops species, barley, and rye. Curr Genet. 2002;41:357–65.

60. Kim DH, Kang JG, Kim BD. Isolation and characterization of the cytoplasmic male sterility-associated orf456 gene of chili pepper (Capsicum annuum L.). Plant Mol Biol. 2007;63:519–32.

61. Arrieta-Montiel MP, Shedge V, Davila J, Christensen AC, Mackenzie SA. Diversity of the arabidopsis mitochondrial genome occurs via nuclear-controlled recombination activity. Genetics. 2009;183:1261–8.

62. Gualberto JM, Mileshina D, Wallet C, Niazi AK, Weber-Lotfi F, Dietrich A. The plant mitochondrial genome: Dynamics and maintenance. Biochimie. 2014;100:107–20.

63. Lupold DS, Caoile a G, Stern DB. Genomic context influences the activity of maize mitochondrial cox2 promoters. Proc Natl Acad Sci U S A. 1999;96:11670–5. doi:10.1073/pnas.96.20.11670.

64. Matera JT, Monroe J, Smelser W, Gabay-Laughnan S, Newton KJ. Unique changes in mitochondrial genomes associated with reversions of S-type cytoplasmic male sterility in Maizemar. PLoS One. 2011;6: e23405.

65. Sloan DB, MacQueen AH, Alverson AJ, Palmer JD, Taylor DR. Extensive loss of RNA editing sites in rapidly evolving silene mitochondrial genomes: Selection vs. retroprocessing as the driving force. Genetics. 2010;185:1369–80.

66. Schmitz-Linneweber C, Kushnir S, Babiychuk E, Poltnigg P, Herrmann RG, Maier RM. Pigment Deficiency in Nightshade/Tobacco Cybrids Is Caused by the Failure to Edit the Plastid ATPase-Subunit mRNA. Plant Cell. 2005;17:1815–28.

67. Gershoni M, Templeton AR, Mishmar D. Mitochondrial bioenergetics as a major motive force of speciation. BioEssays. 2009;31:642–50.

68. Burton RS, Barreto FS. A disproportionate role for mtDNA in Dobzhansky-Muller incompatibilities? Mol Ecol. 2012;21:4942–57.

69. Sloan DB, Havird JC, Sharbrough J. The on-again, off-again relationship between mitochondrial genomes and species boundaries. Mol Ecol. 2017;26:2212–36.

70. Gagnaire PA, Normandeau E, Bernatchez L. Comparative genomics reveals adaptive protein evolution and a possible cytonuclear incompatibility between European and American Eels. Mol Biol Evol. 2012;29:2909–19.

71. Barreto FS, Burton RS. Evidence for compensatory evolution of ribosomal proteins in response to rapid divergence of mitochondrial rRNA. Mol Biol Evol. 2013;30:310–4.

72. Rockenbach K, Havird JC, Grey Monroe J, Triant DA, Taylor DR, Sloan DB. Positive selection in rapidly evolving plastid-nuclear enzyme complexes. Genetics. 2016;204:1507–22.

73. Sloan DB, Triant DA, Wu M, Taylor DR. Cytonuclear interactions and relaxed selection accelerate sequence evolution in organelle ribosomes. Mol Biol Evol. 2014;31:673–82.

74. Edera AA, Gandini CL, Sanchez-Puerta MV. Towards a comprehensive picture of C-to-U RNA editing sites in angiosperm mitochondria. Plant Mol Biol. 2018;97:215–31. doi:10.1007/s11103-018-0734-9.

75. Marker C, Zemann A, Terhörst T, Kiefmann M, Kastenmayer JP, Green P, et al. Experimental RNomics: Identification of 140 candidates for small non-messenger RNAs in the plant Arabidopsis thaliana. Curr Biol. 2002;12:2002–13.

76. Grimes BT, Sisay AK, Carroll HD, Cahoon AB. Deep sequencing of the tobacco mitochondrial transcriptome reveals expressed ORFs and numerous editing sites outside coding regions. BMC Genomics. 2014;15:31.

77. Dietrich A, Wallet C, Iqbal RK, Gualberto JM, Lotfi F. Organellar non-coding RNAs: Emerging regulation mechanisms. Biochimie. 2015;117:48–62.

78. Lowe TM, Eddy SR. tRNAscan-SE: A program for inproved detection of transfer RNA genes in genomic sequence. Nucleic Acids Res. 1997;25:955–64.

79. Lohse M, Drechsel O, Kahlau S, Bock R. OrganellarGenomeDRAW-a suite of tools for generating physical maps of plastid and mitochondrial genomes and visualizing expression data sets. Nucleic Acids Res. 2013;41:W575–81.

80. Krzywinski M et al. Circos: an Information Aesthetic for Comparative Genomics. Genome Res. 2009;19:1639–45. doi:10.1101/gr.092759.109.19.

81. Sloan DB, Alverson AJ, Štorchová H, Palmer JD, Taylor DR. Extensive loss of translational genes in the structurally dynamic mitochondrial genome of the angiosperm Silene latifolia. BMC Evol Biol. 2010;10:274.

82. Edgar RC. MUSCLE: Multiple sequence alignment with high accuracy and high throughput. Nucleic Acids Res. 2004;32:1792–7.

83. Stamatakis A. RAxML version 8: A tool for phylogenetic analysis and post-analysis of large phylogenies. Bioinformatics. 2014;30:1312–3.

84. Librado P, Rozas J. DnaSP v5: A software for comprehensive analysis of DNA polymorphism data. Bioinformatics. 2009;25:1451–2.

85. Štorchová H, Hrdličková R, Chrtek J, Tetera M, Fitze D, Fehrer J. An improved method of DNA isolation from plants collected in the field and conserved in saturated NaCl/CTAB solution. Taxon. 2000;49:79–84.

86. Bolger AM, Lohse M, Usadel B. Trimmomatic: A flexible trimmer for Illumina sequence data. Bioinformatics. 2014;30:2114–20.

87. Wu TD, Nacu S. Fast and SNP-tolerant detection of complex variants and splicing in short reads. Bioinformatics. 2010;26:873–81.

88. McKenna A, Hanna M, Banks E, Sivachenko A, Cibulskis K, Kernytsky A et al. The Genome Analysis Toolkit: A MapReduce framework for analyzing next-generation DNA sequencing data. Genome Res. 2010;20:1297–303.

89. Wagner GP, Kin K, Lynch VJ. Measurement of mRNA abundance using RNA-seq data: RPKM measure is inconsistent among samples. Theory Biosci. 2012;131:281–5.

90. Quinlan AR, Hall IM. BEDTools: A flexible suite of utilities for comparing genomic features. Bioinformatics. 2010;26:841–2.

